## supplemental material for "The phosphatase PP2C12 is a negative player in LRX-RALF-FER-mediated cell wall integrity sensing"

### Appendix

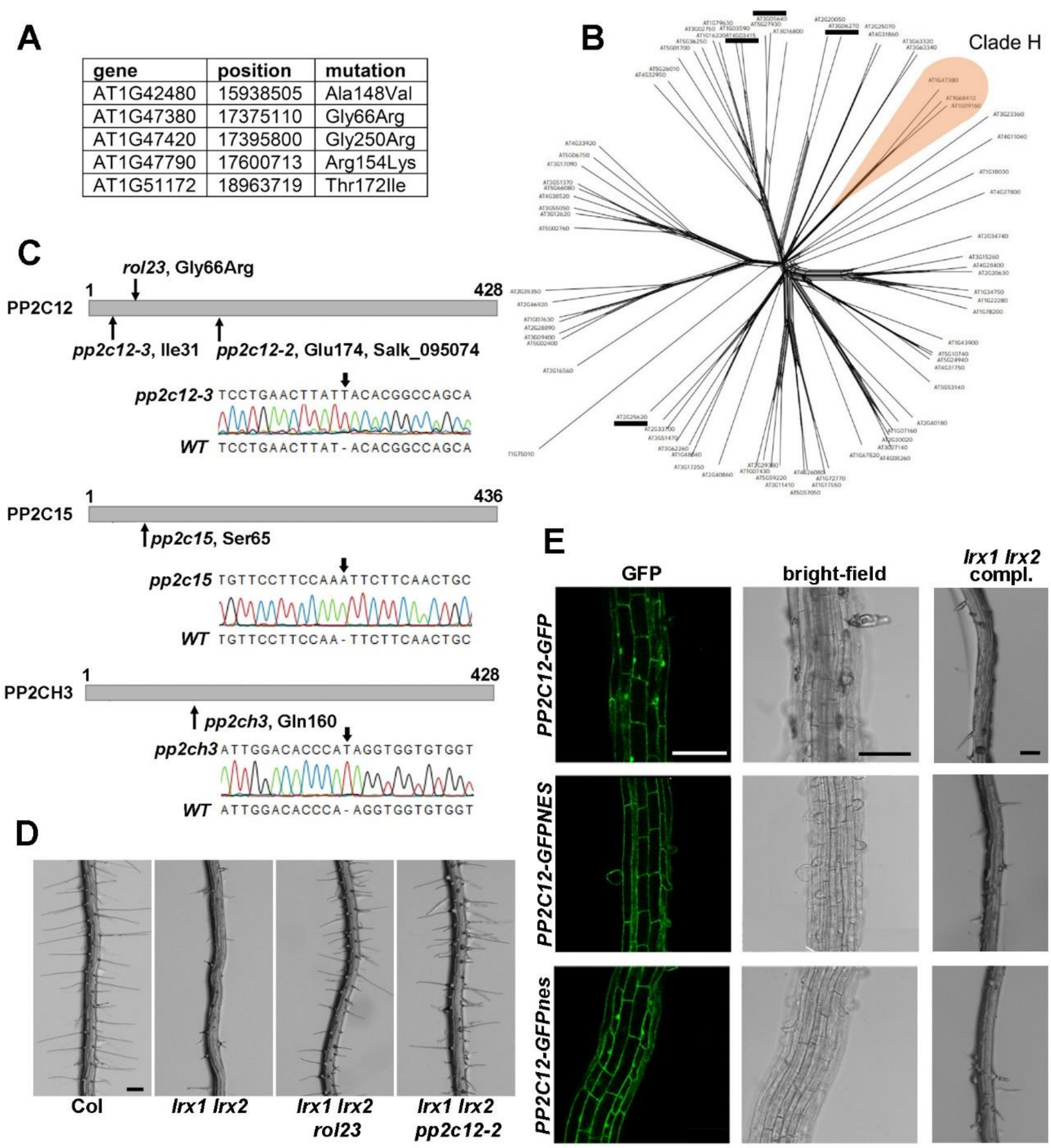

**Appendix Figure S1** ***ROL23*** **codes for PP2C12.**

(A) Whole-genome sequencing revealed several SNPs on chromosome 1 linked to the *rol23* phenotype. (B) Phylogenetic analysis of the PP2C family of phosphatases of *Arabidopsis thaliana*. Clade H PP2Cs are highlighted. PP2Cs of other clades used in this study (see also Table 1) are marked with a black bar. (C) Schematic representation of the clade H PP2C family, the arrows indicating the CRISPR/Cas9-induced 1 bp insertions in all the mutants. (D) The *lrx1 lrx2* double mutant develops an enhanced *lrx1* root hair phenotype, which is also suppressed by *rol23*. (E) *PP2C12::PP2C12-GFP* is ubiquitously expressed in Arabidopsis roots. GFP fluorescence and bright-field images are shown (left and middle panels), and the root hair phenotype of *lrx1 lrx2 rol23* mutants complemented with the constructs (right panels). Size bars = 300 µm.

#### Appendix

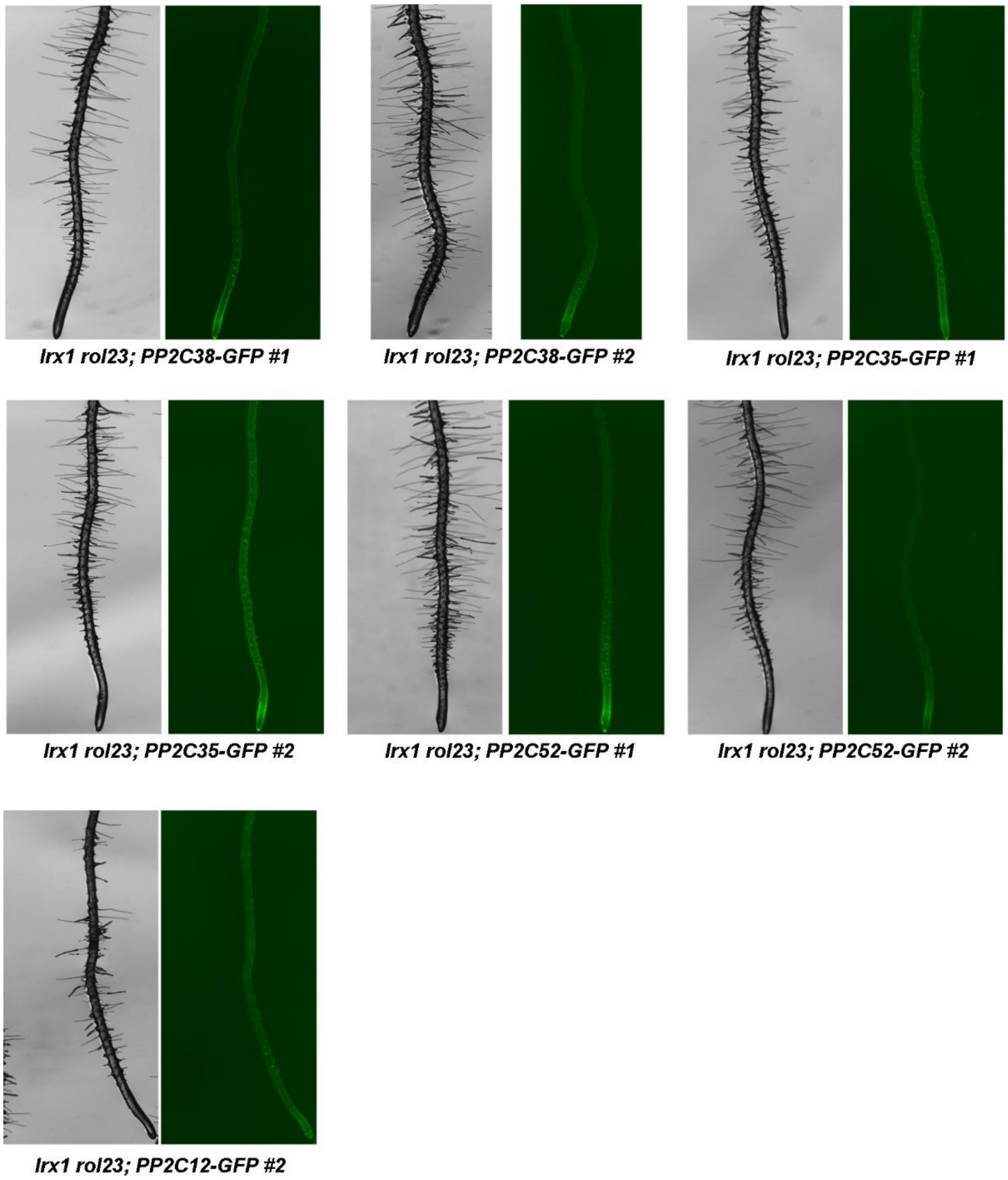

**Appendix Figure S2 Specific function of clade H PP2Cs.**

Representative images of 5-days-old Arabidopsis seedlings grown vertically. Different *PP2C12::PP2Cs-GFP* constructs in the *lrx1 rol23* mutant induce comparable expression levels, i.e. GFP fluorescence, as in *PP2C12::PP2C12-GFP* yet do not complement the *rol23* phenotype. Size bar = 500 µm.

#### Appendix

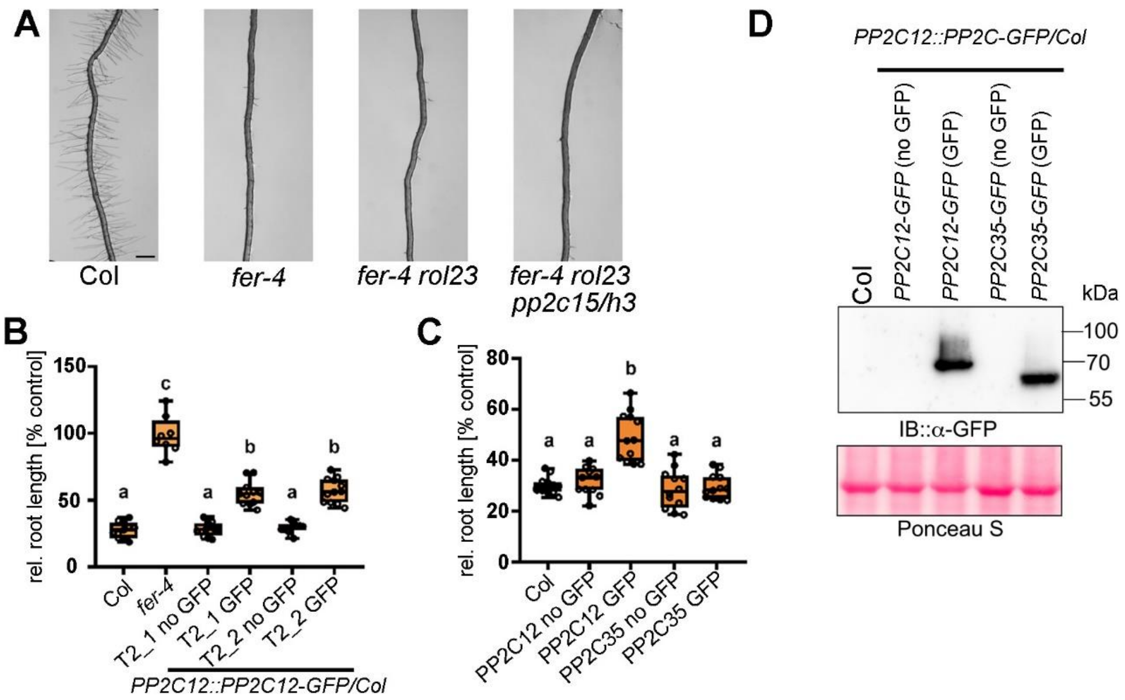

**Appendix Figure S3 PP2C12 affects RALF/FER-related processes.**

(A) Roots of 5-d-old Arabidopsis seedlings grown in a vertical direction. Neither *rol23* nor *rol23 pp2c15 pp2ch3* (*rol23 pp2c15/h3*) have a significant effect on the *fer-4* knock-out root hair phenotype. Size bar = 500 μm. (B,C) Quantification of primary root length of 7-d-old seedlings grown in the absence (mock) or presence of 2 M RALF1 peptide. The primary root length is expressed as relative to mock data for each genotype. Strong expression of *PP2C12::PP2C12-GFP* causes reduced sensitivity to RALF1, as revealed by reduced root growth inhibition. GFP-less seedlings of a population segregating for the transgene show wild type Col-like response to RALF1, but GFP positive lines respond less. This effect is not seen with the non-H-clade *PP2C35*. Different letters indicate significant differences. (one-way ANOVA with Tukey's unequal N-HSD post hoc test, $P < 0.01$). Similar results with at least two independent experiments were obtained. (D) Immunoblot with an anti(α)-GFP antibody on total extracts of seedlings shown in (B) and (C). Blot stained with Ponceau S shows comparable loading. Comparable accumulation of the PP2C12-GFP and PP2C35-GFP were detected, whereas no detectable signals were shown in the GFP-less lines.

#### Appendix

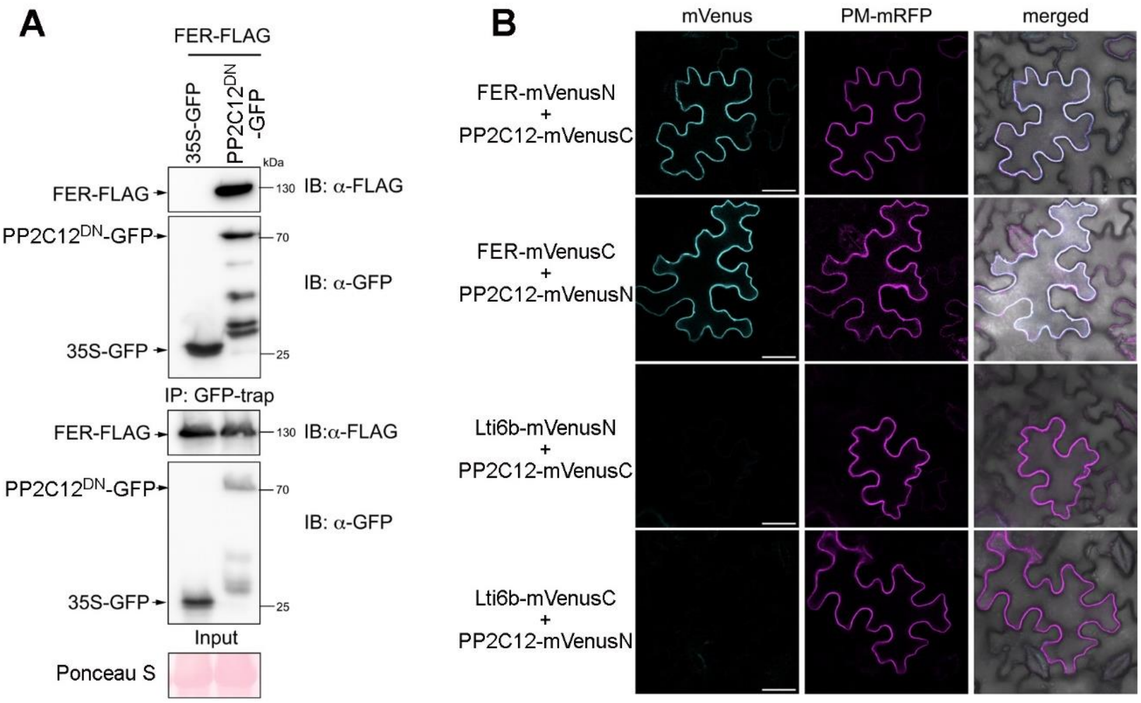

**Appendix Figure S4 PP2C12 interacts with FER at the plasma membrane.**

(A) FER interacts with *PP2C12$^{DN}$* in *N. benthamiana*. Co-IP experiment using overexpressed cytoplasmic GFP as the negative control (opposed to the membrane associated Lti6b-GFP shown in Fig 4). *FER-FLAG* was transiently co-expressed in *N. benthamiana* leaves with either *PP2C12$^{DN}$-GFP* or *GFP*. Immunoprecipitation of PP2C12$^{DN}$-GFP and GFP was done using the anti-GFP trap. FER-FLAG was detected with anti (α)-FLAG antibody when co-expressed with *PP2C12$^{DN}$-GFP* but not *GFP*. Blot stained with Ponceau S shows comparable loading. (B) BiFC assay in *N. benthamiana* as shown in Fig 4 with the addition of the PM marker REM 1.2-mRFP reveals interaction of FER and PP2C12 at the plasma membrane. Size bar= 20 µm.

**Appendix**

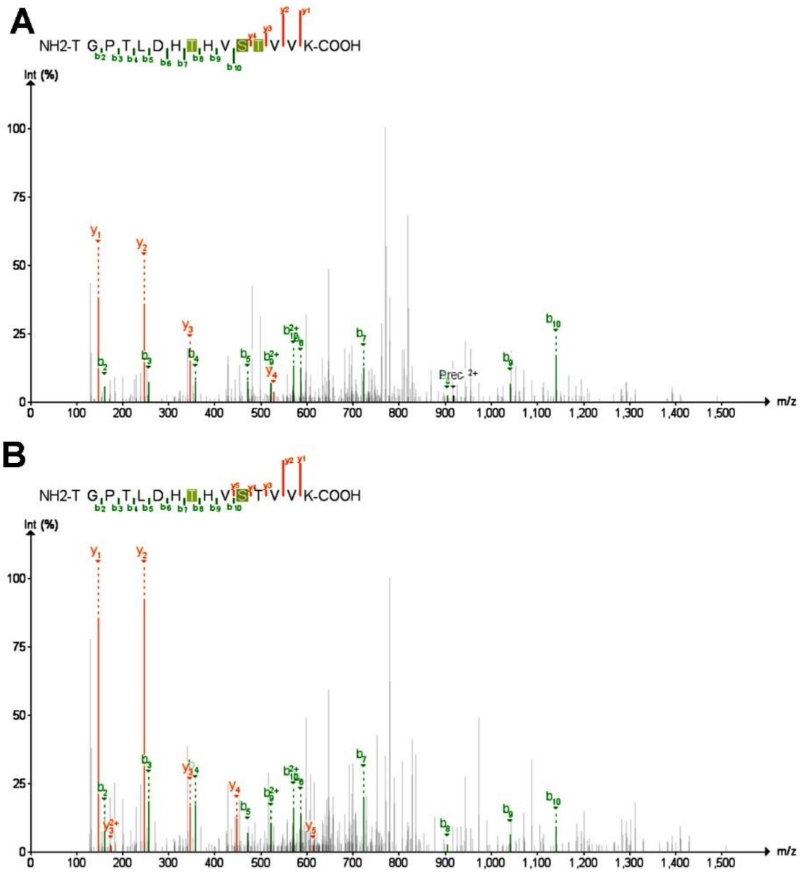

**Appendix Figure S5 MS2 spectra of peptides identifying FER Thr696 as a dephosphorylation target of PP2C12.**

A) The triply phosphorylated peptide carrying pThr692, pSer695, and pThr696 decreased in abundance after treatment with PP2C12. B) The doubly phosphorylated peptide carrying pThr692 and pSer695 increased in abundance after treatment with PP2C12.

**Appendix**

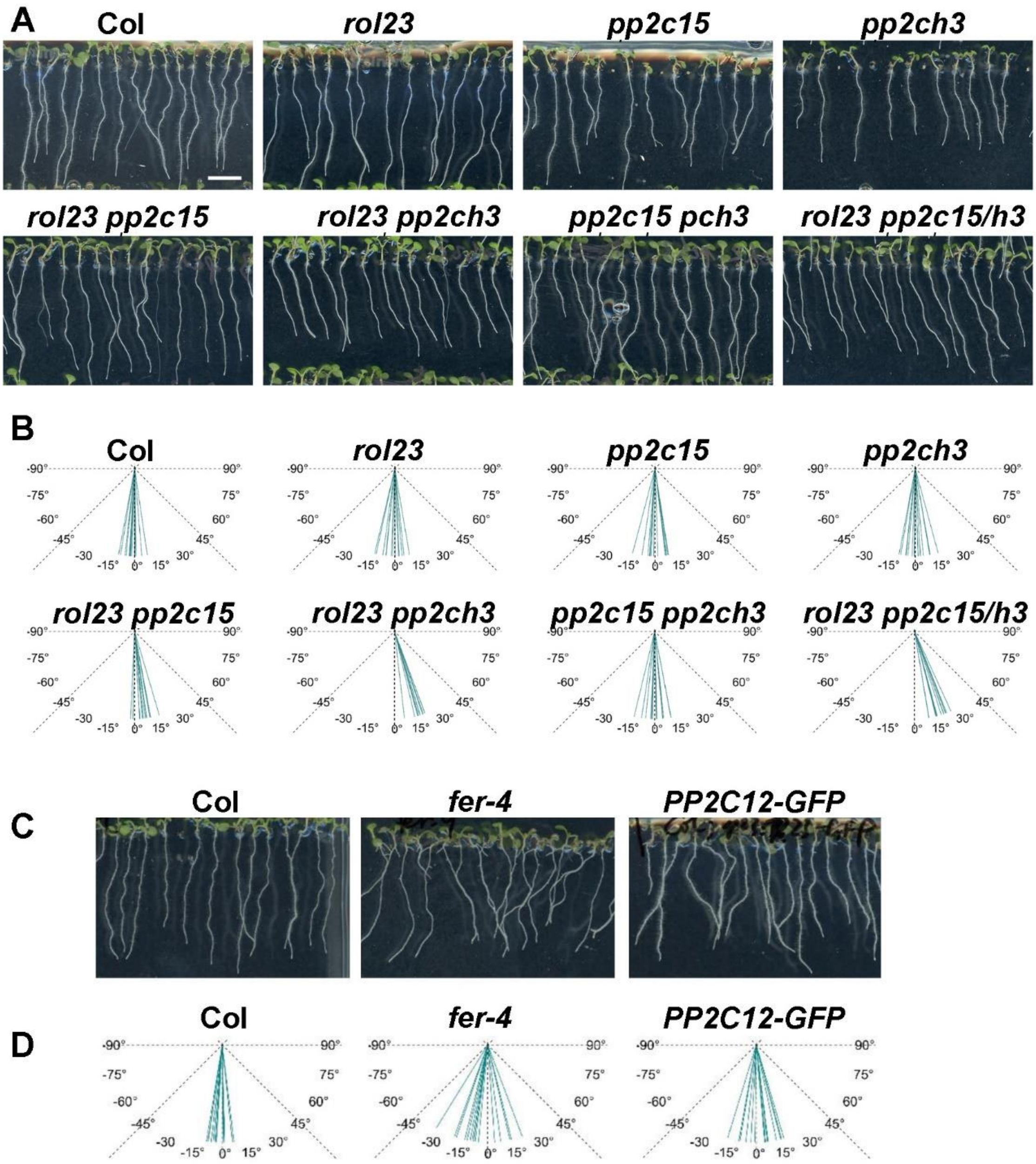

**Appendix Figure S6 Clade H PP2C members redundantly regulate the root skewing.**

(A) Images of 7-d-old Arabidopsis seedlings of Col and clade H *pp2c* mutants grown in a vertical orientation are shown. The primary root skewing is observed in *rol23 pp2c15*, *rol23 pp2ch3*, and *rol23 pp2c15 pp2ch3* (labelled in panel A and B as *rol23 pp2c15/h3*). (B) Quantification of the deviation from the gravitropic vector of seedlings shown in (A). (C) Images of 7-d-old Arabidopsis seedlings of Col, *fer-4* mutant, and a strongly expressing *PP2C12-GFP* line show reduced gravitropism in the latter two lines. (D) Quantification of the deviation from the gravitropic vector of seedlings shown in (C). Bending angles of 15 seedlings per line were measured.

### Appendix

#### Appendix Table S1

**Table S1.** Phosphorylation sites and diagnostic peptides identified on the recombinant FER cytoplasmic domain by LC-MS/MS. Only phosphorylation sites identified by a localization score of >0.75 were included.

| Site | Phosphopeptides(s)[a] | Best Localization Score |
|---|---|---|
| Thr508 | TN**t**TGSYASSLPSNLCR;TN**tt**GSYASSLPSNLCR | 0.9557 |
| Thr509 | TNT**t**GSYASSLPSNLCR;TN**tt**GSYASSLPSNLCR | 0.9479 |
| Ser525 | HF**s**FAEIK;HF**s**FAEIKAATK | 1 |
| Thr533 | AA**t**KNFDE**s**RVLGVGGFGK | 1 |
| Ser539 | AATKNFDE**s**RVLGVGGFGK;AA**t**KNFDE**s**RVLGVGGFGK;NFDE**s**RVLGVGGFGK | 1 |
| Thr559 | VYRGEIDGG**t**TK | 0.9444 |
| Thr560 | VYRGEIDGGT**t**K | 0.823 |
| Ser571 | GNPM**s**EQGVHEFQTEIEMLSK | 0.892 |
| Thr580 | GNPMSEQGVHEFQ**t**EIEMLSK;GNPMSEQGVHEFQ**t**EIEML**s**KLR | 0.9342 |
| Ser586 | GNPMSEQGVHEFQtEIEML**s**KLR | 0.9337 |
| Thr625 | EHLYK**t**QNPSLPWK | 0.8804 |
| Ser629 | TQNP**s**LPWK | 0.9522 |
| Tyr648 | GLH**y**LHTGAKHTIIHR | 0.8224 |
| Thr651 | GLHYLH**t**GAKHTIIHR | 0.9234 |
| Thr656 | GLHYLHTGAKH**t**IIHR | 0.7871 |
| Thr664 | DVK**t**TNILLDEK | 0.9574 |
| Thr665 | DVKT**t**NILLDEK;DVKT**t**NILLDEKWVAK | 0.9391 |
| Ser683 | VSDFGL**s**K;VSDFGL**s**KTGPTLDHTHVSTVVK | 0.9511 |
| Thr685 | VSDFGLSK**t**GP**t**LDHTHVSTVVK | 0.8953 |
| Thr688 | TGP**t**LDHTHVSTVVK;VSDFGLSKTGP**t**LDHtHVSTVVK;VSDFGLSK**t**GP**t**LDHTHVSTVVK | 0.8953 |
| Thr692 | TGPTLDH**t**HVSTVVK;TGPTLDH**t**HV**s**TVVK;TGPTLDH**t**HV**st**VVK;TGPTLDH**t**HV**st**VVKGSFGYLDPEYFR;VSDFGLSKTGP**t**LDH**t**HVSTVVK | 0.9601 |
| Ser695 | TGPTLDHTHV**s**TVVK;TGPTLDHTHV**st**VVKGSFGYLDPEYFR;TGPTLDHTHV**st**VVKG**s**FGYLDPEYFR;TGPTLDH**t**HV**s**TVVK;TGPTLDH**t**HV**st**VVK;TGPTLDH**t**HV**st**VVKGSFGYLDPEYFR | 0.9602 |
| Thr696[b] | TGPTLDHTHVS**t**VVKGSFGYLDPEYFR;TGPTLDHTHV**st**VVKGSFGYLDPEYFR;TGPTLDHTHV**st**VVKG**s**FGYLDPEYFR;TGPTLDH**t**HV**st**VVK;TGPTLDH**t**HV**st**VVKGSFGYLDPEYFR | 0.9602 |
| Ser701 | G**s**FGYLDPEYFR;TGPTLDHTHVSTVVKG**s**FGYLDPEYFR;TGPTLDHTHV**st**VVKG**s**FGYLDPEYFR | 0.963 |
| Ser748 | EQV**s**LAEWAPYCYK;EQV**s**LAEWAPYCYKK | 0.9652 |
| Thr785 | FAE**t**AMK | 1 |
| Ser858 | NDKSSDVYEGNVTD**s**R | 0.7828 |
| Ser866 | SSGIDM**s**IGGR | 0.9408 |

a, Phosphorylated residues are indicated in bold. Some peptides appear in multiple rows where doubly and triply phosphorylated peptides were identified.

b, Dephosphorylated by ROL23.

### Appendix

#### Appendix Table S2

Primers used

| | | |
|---|---|---|
| Primers used for cloning | | |
| ROL23PromF | ATGACTAGTGTGTTAATCTTCTTTGACACATC | |
| ROL23PromR | TGAGGCGCGCCCTTCCTCTAAGCTGCGTCTAG | |
| ROL23CDSF | GGCGCGCCATGTCAACAAAGGGAGAACATC | |
| ROL23CDSR | GGCGCGCCTGTCGCTTTCACTACTATGTC | |
| EGR_F | GACGCAGCTTAGAGGAAGGGCGCGCATGGGACATTTCTCTTCCATG | |
| EGR_R | CTTCTCCTTTACTCATGGCGCGCCATAGAGATGGCGACGACGATG | |
| PP2C28_F | GACGCAGCTTAGAGGAAGGGCGCGCATGGTATCATCGGCAACTATATTG | |
| PP2C28_R | CTTCTCCTTTACTCATGGCGCGCCAAGTAGAAGGTCCAGCTAAATC | |
| PP2C52_F | GACGCAGCTTAGAGGAAGGGCGCGCATGGGGGGTTGTGTGTCGACTAG | |
| PP2C52_R | CTTCTCCTTTACTCATGGCGCGCCAAGTCTTCGATTTCTCTTCAG | |
| PP2C35_F | GACGCAGCTTAGAGGAAGGGCGCGCATGGGTTGTGTTCAATGCAAATG | |
| PP2C35_R | CTTCTCCTTTACTCATGGCGCGCCATTCATTAGACAGCTTTTTAATC | |
| PP2C15_F | TCAGGCGCGCCATGGCGTCTAGAGAAGGAAAG | |
| PP2C15_R | TCAGGCGCGCCAGATGACTTTAACTCCACTTG | |
| PP2CH3_F | GTTCCAGCCAATGCTGAAGGCGCGCATGAGCGTGTCAAAAGCATCG | |
| PP2CH3_R | CTTCTCCTTTACTCATGGCGCGCCAAGTAGTCACTGAACCTTTGC | |
| GFPNESnes_F | CATTACCTGTCCACACAATCTG | |
| NES_R | GATCGGGGAAATTCGAGCTCTCACTTGTTAATATCAAGTCCAGCCAACTTAAGAGC | |
| nes_R | CTCCAGCTGCCTTAAGAGCAAGCTCGTTGTGGTGGTGGTGGTGGTGT | |
| pET28a(+)-FER CD_F | CGCGGATCCGAATTCGCTTACCGCAGACGTAAGCG | |
| pET28a(+)-FER CD_R | CGCAAGCTTGTCGACCTAACGTCCCTTTGGATTCATGA | |
| pMALc4e-MBP_ PP2C12_F | ATCCTCTAGAGTCGACATGTCAACAAAGGGAGAACATC | |
| pMALc4e-MBP_ PP2C12_R | GGCCAGTGCCAAGCTTCTAGTCGCTTTCACTACTATGT | |
| pDEST-Cterm tag-FER FL_F | GGGGACAAGTTTGTACAAAAAAGCAGGCTATGAAGATCACAGAGGGACG | |
| pDEST-Cterm tag-FER FL_R | GGGGACCACTTTGTACAAGAAAGCTGGGTCACGTCCCTTTGGATTCATGA | |
| pDEST-Nterm tag-PP2C12_F | GGGGACAAGTTTGTACAAAAAAGCAGGCTGGATGTCAACAAAGGGAGAACA | |
| pDEST-Nterm tag-PP2C12_R | GGGGACCACTTTGTACAAGAAAGCTGGGTCTAGTCGCTTTCACTACTAT | |
| pDEST-Cterm tag-lti1.3_F | GGGGACAAGTTTGTACAAAAAAGCAGGCTATGAGTACAGCCACTTTCGTAG | |
| pDEST-Cterm tag-lti1.3_R | GGGGACCACTTTGTACAAGAAAGCTGGGTCCTTGGTGATGATATAAAGAGCG | |
| *rol23* CAPS marker | | |
| ROL23_F1 | GACTAACTTGGATTTCAAGTTTCAG | |
| SG1 | AGAGAAAGAAGGCAAATAATAGGCC | mutated bases to introduce StuI site in *rol23* |
| Crispr genotyping | | |
| *pp2c12-3* | | |
| ROL23_F1 | GACTAACTTGGATTTCAAGTTTCAG | |
| ROL23S182_R | CTTCACCACCACTTGCAGTGAC | |
| *pp2c15* | | |
| 410_F5 | CATCAGAGTACTAAGTTAGTTTC | |
| 410_R1 | CTAGGAAGAGCACTAATAACATG | |
| *pp2ch3* | | |
| 160_F1 | CTAAGGAACATCTTCTGGAAAAC | |
| 160_R1 | TTGCATTGGCATAATCAAAGCAC | |
| *Crispr/CAS9* gRNAs | | |
| ROL23 gRNA_2F | attgGACAATCCTGAACTTATACA | |
| ROL23 gRNA_2R | aaacTGTATAAGTTCAGGATTGTC | |
| PP2C15 gRNA_1F | attgAAAGCAGTTGAAGAATTGGA | |
| PP2C15 gRNA_1R | aaacTCCAATTCTTCAACTGCTTT | |
| PP2CH3 gRNA_1F | attgTGCATATTGGACACCCAAGG | |
| PP2CH3 gRNA_1R | aaacCCTTGGGTGTCCAATATGCA | |
